# Meteorin alleviates Paclitaxel-induced peripheral neuropathy in mice

**DOI:** 10.1101/2022.09.13.507857

**Authors:** Ishwarya Sankaranarayanan, Diana Tavares-Ferreira, Lucy He, Moeno Kume, Juliet Mwirigi, Torsten M. Madsen, Kenneth A. Petersen, Gordon Munro, Theodore J. Price

**Affiliations:** Pain Neurobiology Research Group, Department of Neuroscience, Center for Advanced Pain Studies, School of Behavioural and Brain Sciences, University of Texas at Dallas; Richardson, TX, USA; Hoba Therapeutics ApS, Copenhagen, Denmark

**Keywords:** Meteorin, CIPN, Paclitaxel, Satellite glial cells, IENF, Neuropathic Pain, Allodynia

## Abstract

Chemotherapy-induced peripheral neuropathy (CIPN) is a challenging condition to treat, and arises due to severe, dose-limiting toxicity of chemotherapeutic drugs such as paclitaxel. This often results in debilitating sensory and motor deficits that are not effectively prevented or alleviated by existing therapeutic interventions. Recent studies have demonstrated the therapeutic effects of Meteorin, a neurotrophic factor, in reversing neuropathic pain in rodent models of peripheral nerve injury induced by physical trauma. Here, we sought to investigate the potential antinociceptive effects of recombinant mouse Meteorin (rmMeteorin) using a paclitaxel-induced peripheral neuropathy model in male and female mice. Paclitaxel treatment (4 x 4mg/kg, i.p.) induced hind paw mechanical hypersensitivity by day 8 after treatment. Thereafter, in a reversal dosing paradigm, five repeated injections of rmMeteorin (0.5 and 1.8mg/kg s.c. respectively) administered over 9 days produced a significant and long-lasting attenuation of mechanical hypersensitivity in both sexes. Additionally, administration of rmMeteorin (0.5 and 1.8mg/kg), initiated before and during paclitaxel treatment (prevention dosing paradigm), blocked the establishment of hind paw mechanical hypersensitivity. Repeated systemic administration of rmMeteorin in both dosing paradigms decreased histochemical signs of satellite glial cell reactivity as measured by glutamine synthetase and connexin43 protein expression in the DRG. Additionally, in the prevention administration paradigm rmMeteorin had a protective effect against paclitaxel-induced loss of intraepidermal nerve fibers. Our findings indicate that rmMeteorin has a robust and sustained antinociceptive effect in the paclitaxel-induced peripheral neuropathy model and the development of recombinant human Meteorin could be a novel and effective therapeutic for CIPN treatment.

**Highlights:**

- Meteorin produces an antinociceptive effect in both male and female mice treated with paclitaxel.
- Satellite glial cell reactivity induced by paclitaxel treatment is reversed by Meteorin.
- Retraction of intraepidermal nerve fibre (IENF) is blocked by Meteorin treatment in paclitaxel treated mice.
- Findings suggest a disease modifying effect of Meteorin in the mouse model of paclitaxel-induced peripheral neuropathy.

## 1. Introduction

Chemotherapy-induced peripheral neuropathy (CIPN) arises due to neurotoxicity produced by chemotherapy drugs such as paclitaxel and oxaliplatin. The lack of a blood-nerve barrier in the dorsal root ganglion (DRG) allows for accumulation of these chemotherapy drugs resulting in neuronal hyperexcitability and neuropathic pain (McWhinney et al., 2009; Gornstein and Schwarz, 2014; Megat et al., 2019). Characteristics of CIPN include numbness, tingling, hypersensitivity to cold and mechanical stimuli, and burning sensations in the hands and feet (Krishnan et al., 2005; Boyette-Davis et al., 2013). It affects around 30-40% of cancer patients and can continue to persist even after cessation of treatment (Seretny et al., 2014). Currently, there are no FDA-approved drugs for the treatment of CIPN. Paclitaxel which can induce neuropathy as a dose-limiting side effect accumulates in the DRG and causes direct neurotoxic effects such as axonal impairment and neuronal mitochondrial damage (Boyette-Davis et al., 2011; Malacrida et al., 2019). Approximately 25% of patients with paclitaxel induced neuropathy discontinue their treatment or reduce their dosage primarily due to pain (Salgado et al., 2020). Hence, identifying a novel drug for the treatment of paclitaxel neuropathy would be hugely beneficial and be expected to improve both survival probability the quality of life for cancer patients.

Neurotrophic factors (NTF) promote growth, survival, differentiation, and maintenance of cells of the nervous system. These factors can promote a therapeutic effect in many neurodegenerative disorders and also in neuropathic pain (Sah et al., 2005; Ossipov, 2011). Glial-derived neurotrophic factor (GDNF), neurturin (NTRN), and artemin belong to a transforming growth factor β superfamily subgroup. Glial-derived neurotrophic factor was the first glial cell-derived NTF to show neuroprotection, and it signals through a GPI linked to the RET tyrosine kinase (Treanor et al., 1996). Artemin has been shown to support the survival of sensory neurons in culture through interacting with a GFR-a3 receptor primarily expressed in the peripheral nervous system (Gardell et al., 2003). Glial-derived neurotrophic factor (GDNF) administration also reverses mechanical hypersensitivity in rats with spinal nerve ligation (SNL) (Baloh et al., 1998; Boucher et al., 2000). These and other studies on growth factors for pain treatment suggest great potential for antinociceptive effects, and even disease modification, but no treatments in this class have so far been approved for pain relief (Pezet and McMahon, 2006; Lewin et al., 2014). Treatments sequestering growth factors have been tested in clinical trials for pain. Anti-NGF therapeutics achieved robust superiority over placebo in many clinical trials (Lane et al., 2010; Katz et al., 2011; Brown et al., 2012, 2013; Kivitz et al., 2013), but these therapeutics have not been approved for human use due to side-effects.

Meteorin is a neurotrophic factor which produces long-lasting antinociceptive effects in rodent models of peripheral nerve injury induced by physical trauma (Jorgensen et al., 2012; Xie et al., 2019a). It is expressed in the central and peripheral nervous system, with high expression levels in the DRG in humans and mice, where it is likely expressed by DRG neurons and satellite glial cells based on single cell sequencing experiments (Ray et al., 2018; Zeisel et al., 2018; Tavares-Ferreira et al., 2022). Meteorin promotes neurite outgrowth in neurons, including DRG neurons where it has an effect on small and intermediate-sized neurons, which are presumably nociceptors (Nishino et al., 2004; Jørgensen et al., 2009; Jørgensen et al., 2012). Meteorin also promotes glial cell differentiation via the activation of the JAK-STAT pathway, but the underlying receptor-mediated mechanisms leading to these effects are still unclear (Lee et al., 2010). Two previous studies have demonstrated the therapeutic effects of Meteorin in reversing neuropathic pain induced in the sciatic nerve injury in rats (Jørgensen et al., 2012; Xie et al., 2019b). The goal of this study was to test the hypothesis that recombinant mouse Meteorin (rmMeteorin) would be able to prevent and reverse paclitaxel-induced peripheral neuropathic pain in male and female mice.

In support of our hypothesis, we demonstrate that systemic administration of rmMeteorin (0.5mg/kg and 1.8mg/kg) is efficacious in prevention and reversal paradigms in the mouse paclitaxel-induced neuropathy model with mechanical hypersensitivity as the primary endpoint. Administration of rmMeteorin also reduced the expression of satellite cell gliosis.

Finally, rmMeteorin partially halted intraepidermal nerve fiber (IENF) loss in the prevention dosing paradigm. Overall, our findings demonstrate that the development of recombinant human Meteorin could be a novel and effective therapeutic for CIPN treatment.

## 2. Material and Methods

### 2.1 Animals

ICR (CD-1) male and female mice were purchased from Envigo and maintained at the animal facility at the University of Texas at Dallas. Mice were group-housed (4 maximum) in cages with bedding material provided for enrichment with food and water available *ad libitum* in a 12:12 h light-dark cycle. Room temperature was maintained at 21-22°C. Behavioral experiments were performed on mice ranging between 8-12 weeks old at the start of the experiment. All procedures were approved by the Institution Animal Care and Use Committee (IACUC) at the University of Texas at Dallas. Both male and female mice were used for behavioral testing in the reversal paradigm and only female mice were used for behavioral testing in the prevention paradigm.

### 2.2 Injections

Administered drug was prepared by dissolving Paclitaxel in 50% EL Kolliphor (Sigma-Aldrich) and 50% ethanol and further diluted in sterile Dulbecco’s phosphate buffered saline (DBPS; Thermo Scientific). Mice were administrated 4 mg/kg every-other-day for four days for a cumulative dosage of 16 mg/kg or vehicle control of 50% EL Kolliphor −50% ethanol diluted in DPBS. Paclitaxel was injected interperitoneally (i.p) on days two, four, six and eight using a 27-gauge needle (Lees et al.). Starting on day 10, in a reversal paradigm, five subcutaneous injections of recombinant mouse Meteorin (rmMeteorin, R & D Systems #3475) either 0.5mg/kg or 1.8mg/kg or vehicle control (DPBS) was administered every other day for ten days using an Insulin syringe (Lees et al.) with a 30 gauge needle. For the prevention paradigm, subcutaneous rmMeteorin 0.5 or 1.8mg/kg or vehicle was administered before and then given intermittently between paclitaxel treatments on days 1,3,5,7 and 9. The investigator was blinded to treatment during all the injections.

### 2.3 Mechanical withdrawal thresholds

Behavioral testing was performed after habituating mice for 2 hrs to clear acrylic behavioral chambers before beginning mechanical testing. Mechanical hypersensitivity was tested at baseline and every other day, days with no injections or as indicated in the figures. Mechanical paw withdrawal threshold was tested with the up-down method (Chaplan et al., 1994) using calibrated von Frey filaments (Stoelting) applied to the mid plantar surface of the left hind paw. A positive response was comprised of licking or immediate flicking of the hind paw upon application of the filament. The investigator was blinded on all days of testing.

On day 24, three mice per group were euthanized to harvest tissue for immunohistochemistry. Mechanical paw withdrawal thresholds continued to be assessed in the remaining mice until they had resolved back to a baseline level. Two mice were removed from the male cohort on day nine of the reversal paradigm as they were not hypersensitive after the last paclitaxel treatment. Male behavior testing in the reversal paradigm was halted after day 21 and resumed on day 51 due to a mandatory COVID lockdown in March-April 2020. None of the mice were removed from the female cohorts of both paradigms.

### 2.4 Immunohistochemistry (IHC)

On day 24, three animals per group were anesthetized using 4% isoflurane and euthanized by decapitation. Dorsal root ganglia (DRG) and skin tissues were flash-frozen in Optimum cutting temperature (OCT) medium (Fisher Scientific) on dry ice. Twenty μm sections were cut on a cryostat and mounted onto SuperFrost Plus slides (Thermo Fisher Scientific). The sections were fixed in ice-cold 10% formalin for 15 minutes followed by incubation in an increasing percentage of ethanol 50%, 70%,100% for 5 minutes each. The fixed slides were transferred into blocking solution (10% Normal Goat Serum, 0.3% Triton-X 100 in 0.1 M PB) for 1 hr at room temperature. Sections were incubated in primary antibody diluted in blocking solution for 3 hr at room temperature or 4°C overnight. They were then washed in 0.1 M PB followed by incubation in secondary antibody diluted in blocking solution for 1h at room temperature. The slides were washed again with 0.1 M PB followed by incubation with DAPI diluted in blocking solution for 5 minutes at room temperature. Lastly, the sections were washed in 0.1 M PB and cover-slipped using Prolong Gold Antifade (Thermo Fisher Scientific). Images were taken using an Olympus FluoView 3000 confocal microscope and analyzed using Cellsens (Olympus) software. All IHC images are a representative of a sample size of 3 animals per group.

Image analyses of the DRG sections were performed by drawing a region of interest (ROI) around each individual neuron and the mean grey intensity (MGI) value was calculated in the targeted channel of interest. Average MGI was subtracted from the background fluorescence intensity calculated using the negative control (secondary antibody only without primary antibody) and normalized over the area surrounding each individual neuronal cell body.

Image analysis of IENF density on skin sections was done over the plantar surface of the hind paw (Siau et al., 2006) by calculating the number of fibers crossing the basement membrane over the length of the epidermal membrane per millimeter. Three sections per animal were analyzed with 3 animals per group to calculate the IENF density.

**Table 1:** List of antibodies used for immunohistochemistry:

| Antibodies | Company | Catalog # | Dilution |
| --- | --- | --- | --- |
| Glutamine Synthetase polyclonal antibody | Thermo fisher | Cat # 11307-2-AP | 1:1000 |
| Peripherin | EnCor Biotechnology Inc | Cat # CPCA- Peri | 1:1000 |
| DAPI | Cayman | Item no, 14285 | 1:5000 |
| Connexin 43 | Cell Signaling Technologies | Cat # 3512S | 1:1000 |
| IgG (H+L) Cross-Adsorbed Goat anti-Rabbit, Alexa Fluor® 555 | Fisher Scientific | A21428 | 1:2000 |
| IgY (H+L) Goat anti-Chicken, Alexa Fluor® 647 | Fisher Scientific | A21449 | 1:2000 |

### 2.4 Data and Statistical Analysis

Data analysis was done using Graphpad Prism 8.4.1. Statistical differences between groups were assessed using one-way ANOVA followed by Tukey multiple comparisons. P-values are reported in the figures and figure legends. Behavioral analysis was determined using mixed-effects two-way ANOVA analysis followed by Tukey post hoc tests. The mixed-effect model was used to account for the missing values starting on day 24 when 4 animals per group were euthanized for IHC. Effect sizes were determined by subtracting behavior scores from baseline values. Absolute values were summed from the beginning of rmMeteorin administration (from day 10 for the reversal paradigm and from day 1 for the prevention paradigm) and plotted for each group and compared by one-way ANOVA. All data are represented as mean +/− SEM with p<0.05 considered significant. The sample size and sex are noted in the figures and figure legends.

## 3. Results

### 3.1 Systemic administration of rmMeteorin reverses hindpaw mechanical hypersensitivity in paclitaxel treated female and male mice

Meteorin produces a robust reversal of neuropathic hypersensitivity in various rat models of trauma-induced peripheral nerve injury (Jørgensen et al., 2012; Xie et al., 2019a). We sought to investigate the potential antinociceptive effects of rmMeteorin in paclitaxel-induced peripheral neuropathy in male and female mice. We induced neuropathic pain by administering paclitaxel at 4 mg/kg every other day for 4 days. This was followed by 5 subcutaneous injections of rmMeteorin at 0.5mg/kg or 1.8mg/kg dosage or vehicle (DPBS) given every-other-day for a total of 10 days (Fig 1A). Administration of paclitaxel induced mechanical hypersensitivity in all groups and both sexes of mice by day 9. Thereafter, administration of both doses of rmMeteorin (0.5mg/kg or 1.8mg/kg) produced a robust and sustained reversal of mechanical hypersensitivity in male and female mice (Fig 1B, C). Mechanical paw withdrawal thresholds were tested until all animals in all groups returned to baseline. Effect size was significantly different from vehicle for both doses of rmMeteorin (0.5mg/kg and 1.8mg/kg) demonstrating efficacy in both sexes (Fig 1 D, E).

**Figure 1:**
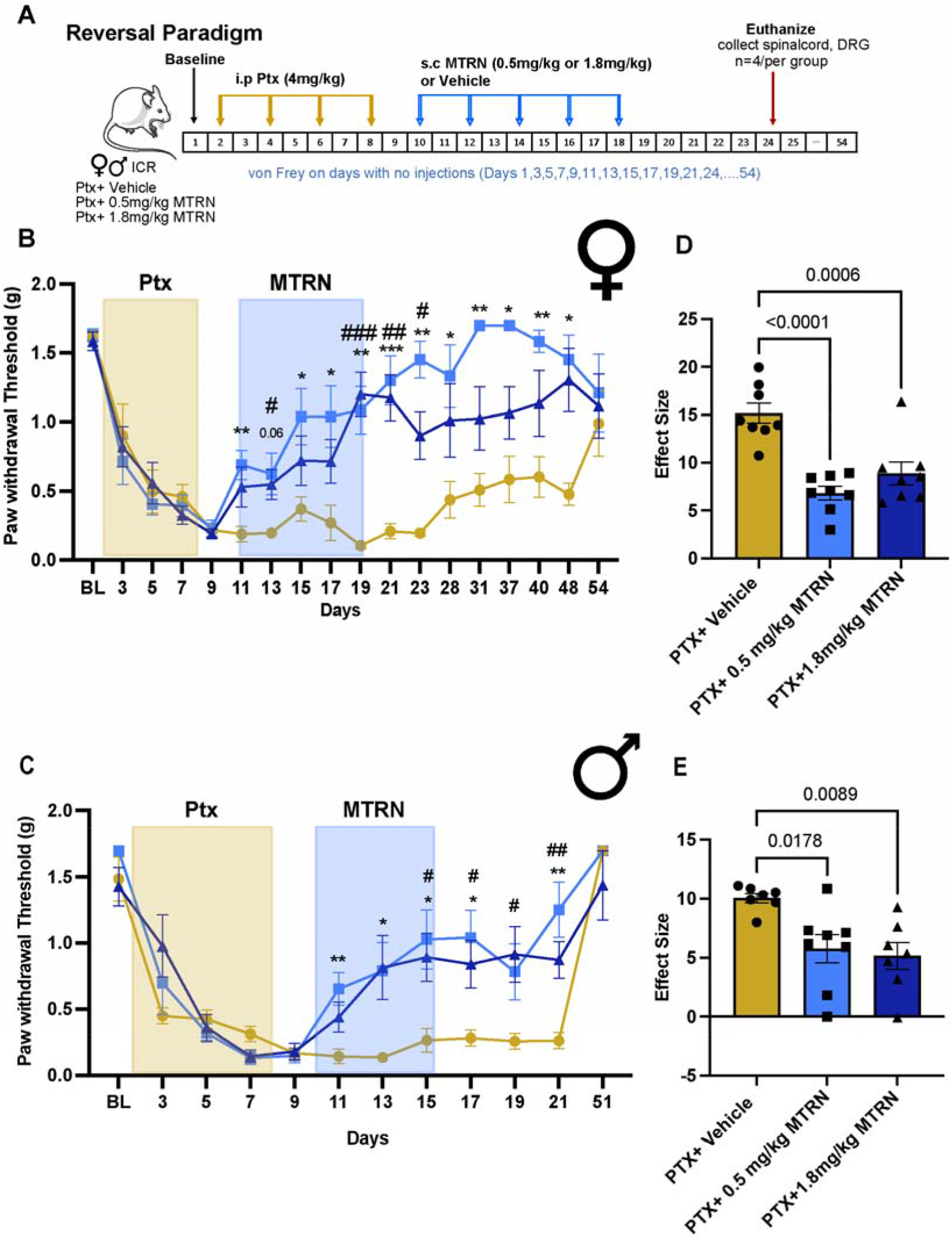
Systemic administration of rmMeteorin reverses hindpaw mechanical hypersensitivity in paclitaxel treated female and male mice. A) A cohort of male and female mice were administered an i.p. injection of 4 mg/kg paclitaxel every other day for a cumulative dose of 16 mg/kg. This was, followed by 5 s.c. injections of 0.5 mg/kg or 1.8 mg/kg of rmMeteorin or Vehicle. Hind paw mechanical thresholds were measured as shown in the figure. B) Mechanical paw withdrawal thresholds in female groups injected with rmMeteorin (0.5 mg/kg and 1.8 mg/kg) displayed reduced mechanical hypersensitivity compared to the paclitaxel (Ptx) + vehicle group. N = 8/group until day 23, N = 4/group until day 54 (Two-way ANOVA-mixed effect analysis, F=4.412, p-value<0.0001, post-hoc Tukey, Ptx + vehicle vs. Ptx + 0.5 mg/kg MTRN, p-value=0.0038 at day 11, Ptx + vehicle vs. Ptx + 1.8 mg/kg MTRN, p-value=0.033 at day 13, Ptx + vehicle vs. Ptx + 0.5 mg/kg MTRN, p-value=0.0335 at day 15, Ptx + vehicle vs. Ptx + 0.5mg/kg MTRN, p-value=0.0292 at day 17, Ptx + vehicle vs. Ptx + 0.5mg/kg MTRN, p-value=0.0018, Ptx + vehicle vs. Ptx + 1.8mg/kg MTRN, p-value=0.0005 at day 19, Ptx + vehicle vs. Ptx + 0.5 mg/kg MTRN, p-value=0.0008, Ptx + vehicle vs. Ptx + 1.8 mg/kg MTRN, p-value=0.0011 at day 21, Ptx + vehicle vs. Ptx + 0.5mg/kg MTRN, p-value= <0.0001, Ptx + vehicle vs. Ptx + 1.8mg/kg MTRN, p-value=0.0107 at day 23, Ptx + vehicle vs. Ptx + 0.5mg/kg MTRN, p-value=0.00436 at day 28, Ptx + vehicle vs. Ptx + 0.5mg/kg MTRN, p-value=0.0042 at day 31, Ptx + vehicle vs. Ptx + 0.5mg/kg MTRN, p-value=0.0139 at day 37, Ptx + vehicle vs. Ptx + 0.5mg/kg MTRN, p-value=0.0058 at day 40, Ptx + vehicle vs. Ptx + 0.5mg/kg MTRN, p-value=0.0145 at day 48. C) Effect size was determined by calculating the cumulative difference between the value for each time point and the baseline value and summed from the beginning of rmMeteorin administration. The effect size difference was significant in the female cohort of mice (Effect size, One-way ANOVA, F=18.91, p-value = <0.0001, post-hoc Tukey, vehicle vs. Ptx + 0.5 mg/kg MTRN, p-value= <0.0001, Ptx + vehicle vs. Ptx + 1.8 mg/kg MTRN, p-value=0.0006). D) rmMeteorin (0.5mg/kg and 1.8mg/kg) administration reversed mechanical withdrawal thresholds in male mice, N = 7-8 until day 21 and N = 3-4 on day 51 (Two-way ANOVA-mixed effect analysis, F=3.349, p-value<0.0001, post-hoc Tukey, Ptx + vehicle vs. Ptx + 0.5 mg/kg MTRN, p-value=0.0096 at day 11, Ptx + vehicle vs. Ptx + 0.5 mg/kg MTRN, p-value=0.0458 at day 13, Ptx + vehicle vs. Ptx + 0.5 mg/kg MTRN, p-value=0.0274, Ptx + vehicle vs. Ptx + 1.8 mg/kg MTRN, p-value=0.0321 at day 15, Ptx + vehicle vs. Ptx + 0.5 mg/kg MTRN, p-value=0.0186, Ptx + vehicle vs. Ptx + 1.8 mg/kg MTRN, p-value=0.0488 at day 17, Ptx + vehicle vs. Ptx + 1.8 mg/kg MTRN, p-value=0.0488 at day 19, Ptx + vehicle vs. Ptx + 0.5 mg/kg MTRN, p-value=0.0044, Ptx + Vehicle vs. Ptx + 1.8 mg/kg MTRN, p-value=0.0093 at day 21. E) Effect size difference in male cohorts of reversal paradigm was significant for both MTRN doses, (One-way ANOVA, F= 6.79, p-value= 0.0.006, post-hoc Tukey, Ptx + vehicle vs. Ptx + 0.5 mg/kg MTRN, p-value=0.0178, Ptx + vehicle vs. Ptx + 1.8 mg/kg MTRN, p-value=0.0089). Data represents mean +/− SEM. Significance represented as Ptx+ Vehicle vs. 0.5 mg/kg MTRN* and Ptx+ Vehicle vs. 1.8 mg/kg MTRN#.

### 3.2 Systemic administration of rmMeteorin reverses satellite cell gliosis caused by paclitaxel treatment

Gliosis of satellite glial cells occurs after chemotherapy treatment causing the cells to change their morphology and release mediators that may contribute to neuronal hyperexcitability associated with neuropathic pain (Liu et al., 2019; Fumagalli et al., 2021).

Therefore, we examined changes in satellite glial cell reactivity within the DRG of the treated cohorts. We assessed immunoreactivity for Glutamine Synthetase (Makker et al., 2017) which labels satellite glial cells that surround the neuronal cell bodies. Paclitaxel administration induced an increase in the expression of GS in both male and female mice (Fig 2A, B). Administration of rmMeteorin (0.5 mg/kg) in the reversal paradigm significantly reduced the expression of GS induced by paclitaxel in both sexes. The 1.8 mg/kg dosage of rmMeteorin also reduced GS expression significantly, albeit only in males (Fig 2 C, D).

**Figure 2:**
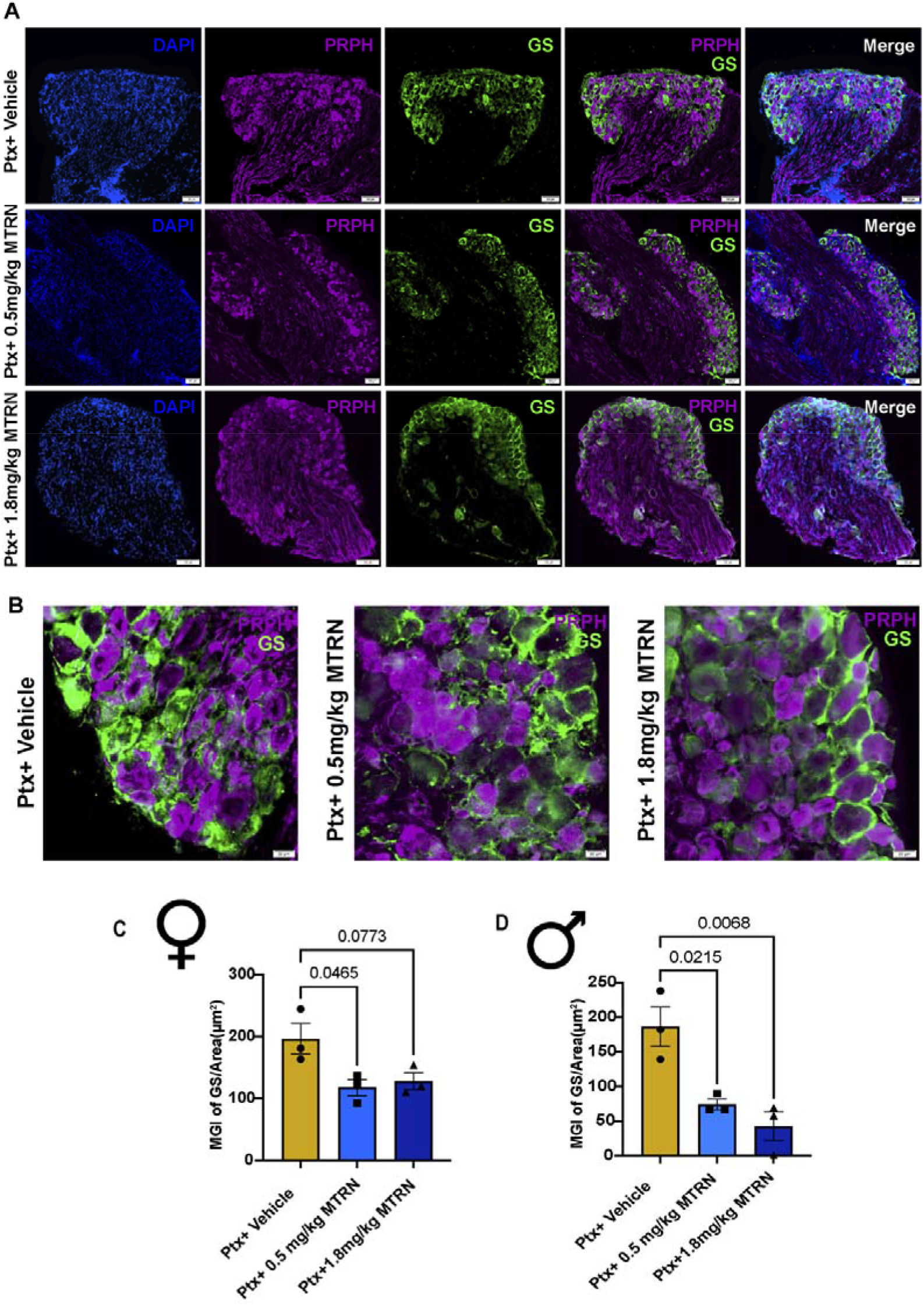
rmMeteorin treatment reverses the satellite cell gliosis in the reversal dosing paradigm of paclitaxel treated mice. A) Representative images show the expression of glutamine synthetase expression in satellite glial cells in green, peripherin (purple) to label neurons, DAPI (blue) to label nuclei. B) Representative images of satellite glial cells at higher magnification show reduced expression of GS in Ptx + 0.5 mg/kg MTRN and Ptx + 1.8 mg/kg MTRN groups compared to Ptx + vehicle. C-D) 0.5 mg/kg MTRN (royal blue) and 1.8 mg/kg MTRN (navy blue) reverses satellite cell gliosis expression compared to Ptx + Vehicle (yellow) in both male and female cohorts of mice in the reversal paradigm (females, One-way ANOVA F=5.779, p-value=0.0399, post-hoc Tukey, vehicle vs. Ptx + 0.5 mg/kg MTRN, p-value= 0.0465, males, One-way ANOVA, F=12.97, p-value=0.0066, vehicle vs. Ptx + 0.5mg/kg MTRN, p-value=0.0215, Ptx + vehicle vs. Ptx + 1.8 mg/kg MTRN, p-value=0.0068). N= 3/group. Data are represented as mean ± SEMs. Scale bar = 100 μm (A), and 20 μm (B).

### 3.3 Recombinant mouse Meteorin reduces connexin43 gap junction expression in mice treated with paclitaxel

Satellite cell gliosis caused by chemotherapy is also associated with an increase in neuronal coupling and expression of gap junctions within DRGs (Warwick and Hanani, 2013). We assessed connexin43 immunolabeling in DRGs of mice treated with paclitaxel with or without rmMeteorin treatment. We observed an increase in connexin43 expression in paclitaxel treated groups in both sexes (Fig 3A, B). Administration of rmMeteorin (0.5 mg/kg) in the reversal paradigm significantly reduced the expression of connexin43 in both males and females. In contrast, administration of 1.8 mg/kg rmMeteorin had a significant effect only in males (Fig 3 C, D).

**Figure 3:**
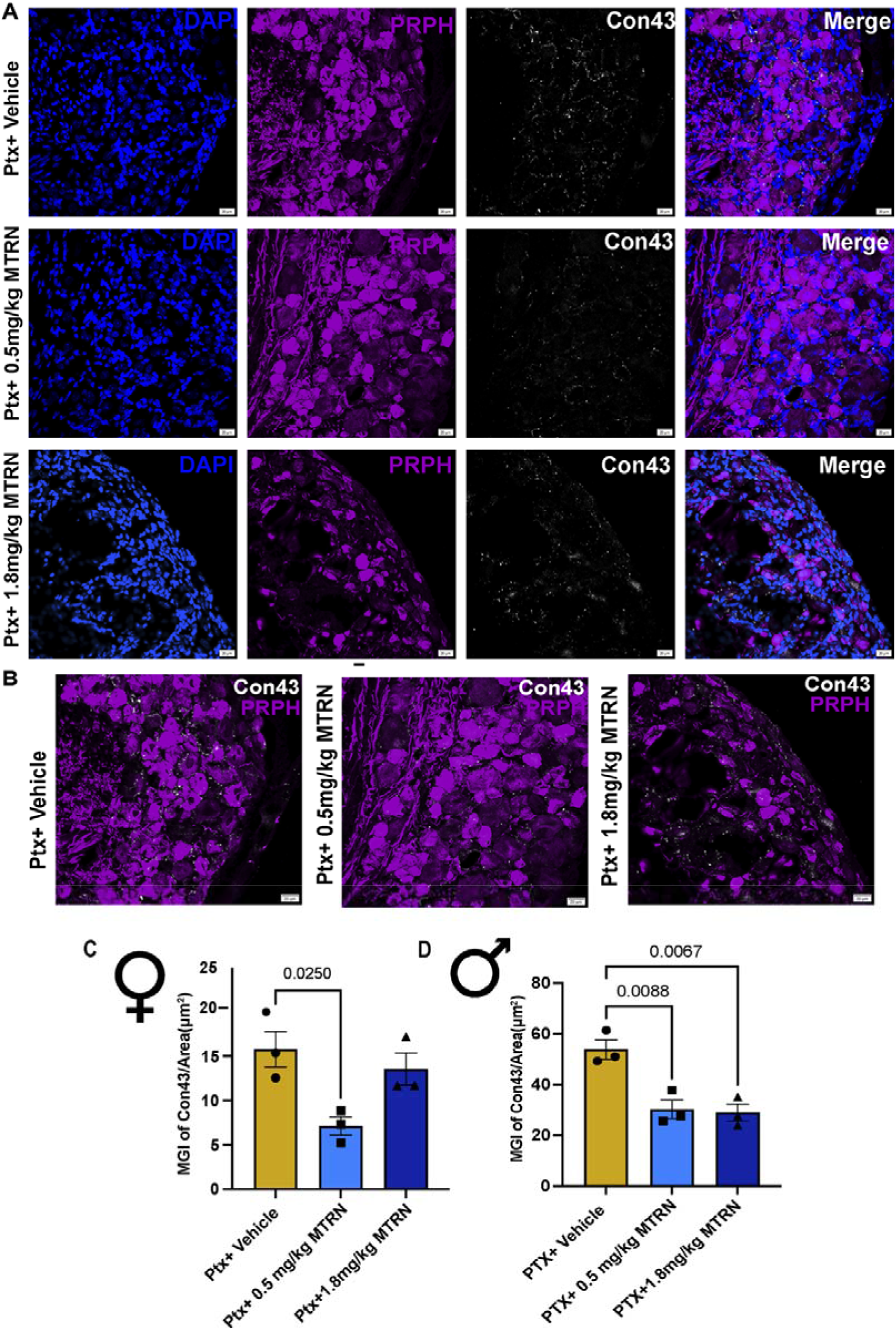
rmMeteorin reduces gap junction expression in mice treated with paclitaxel. A) Representative 20X images of gap junction expression in the DRG showing connexin 43 (white), peripherin (purple) DAPI (blue). Representative 40X overlay images of gap junction expression in DRG of mice using connexin43 (white) with peripherin (purple) in the reversal paradigm groups Ptx + vehicle, Ptx + 0.5 mg/kg MTRN, Ptx + 1.8 mg/kg MTRN. C) Connexin 43 expression in the DRG was significantly reduced in animals treated with rmMeteorin compared to Ptx + vehicle treated mice in both females and males (Females, One-way ANOVA, F=7.170, p-value=0.0257, post-hoc Tukey, vehicle vs. Ptx + 0.5 mg/kg MTRN, p-value=0.025, Ptx + vehicle vs. Ptx + 1.8 mg/kg MTRN, p-value>0.05, Males, One-way ANOVA, F=14.95, p-value=0.0.0047, vehicle vs. Ptx + 0.5 mg/kg MTRN, p-value=0.0088, Ptx + vehicle vs. Ptx + 1.8 mg/kg MTRN, p-value=0.0067) N=3/group. Data are represented as mean ± SEMs. Scale bar = 20 μm.

### 3.4 rmMeteorin blocks development of hind paw mechanical hypersensitivity in the prevention dosing paradigm

Having shown that rmMeteorin could markedly reverse mechanical hypersensitivity when administered after paclitaxel treatment, we sought to explore whether rmMeteorin could block the development of mechanical hypersensitivity in paclitaxel-induced neuropathy using the prevention dosing paradigm in female mice. We used female mice for these experiments because (i) similar effects were seen in both sexes in the interventive experiment and (ii) paclitaxel is reported to cause toxicities more frequently in women (Ozdemir et al., 2018). Accordingly, we injected subcutaneous rmMeteorin 0.5mg/kg, 1.8mg/kg or vehicle starting 1 day before and then on days between paclitaxel treatments (Fig 4A). Here, we observed that both doses of rmMeteorin attenuated the development of paclitaxel induced mechanical hypersensitivity (Fig 4B). Again, the effect of rmMeteorin was both robust and sustained, with the effect size significantly different from vehicle for both doses (Fig 4C).

**Figure 4:**
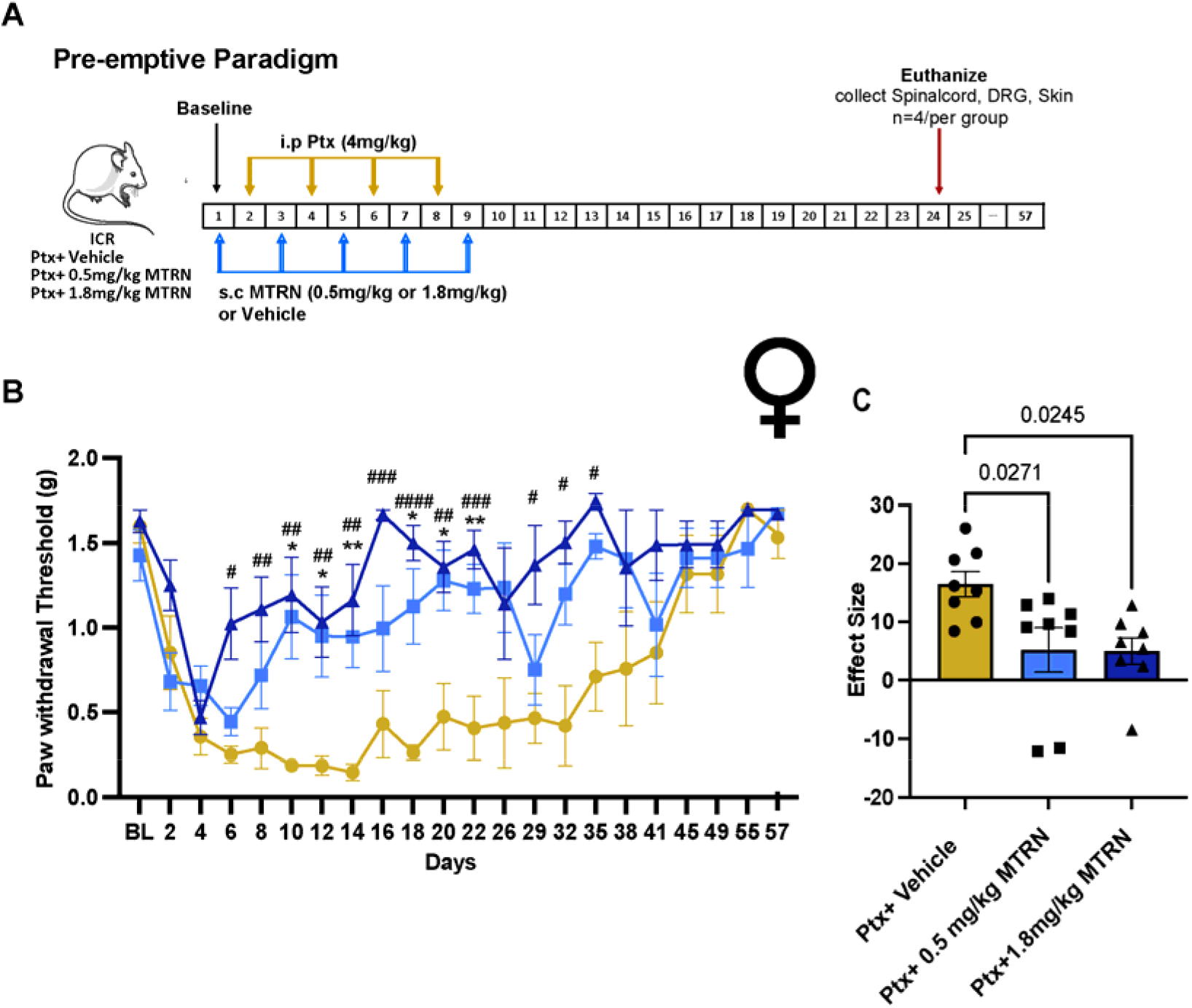
rmMeteorin attenuates development of hind paw mechanical hypersensitivity in the prevention dosing paradigm of paclitaxel treatment. A) Diagram showing the dosing paradigm and testing schedule in female mice. B) 0.5 mg/kg MTRN and 1.8 mg/kg MTRN attenuated development of hind paw mechanical hypersensitivity produced by paclitaxel (Two-way ANOVA mixed effect model, F= 2.425, p-value <0.0001, Tukey multiple comparison, Ptx + vehicle vs. Ptx + 1.8 mg/kg MTRN, p-value= 0.0157 on day 6, Ptx + vehicle vs. Ptx + 1.8 mg/kg MTRN, p-value= 0.0069 on day 8, vehicle vs. Ptx + 0.5 mg/kg MTRN, p-value= 0.0192, Ptx + vehicle vs. Ptx +1.8 mg/kg MTRN, p-value=0.0051 on day 10, vehicle vs. Ptx + 0.5 mg/kg MTRN, p-value= 0.0327, Ptx + vehicle vs. Ptx + 1.8 mg/kg MTRN, p-value=0.0083 on day 12, vehicle vs. Ptx + 0.5 mg/kg MTRN, p-value= 0.0055, Ptx + vehicle vs. Ptx + 1.8 mg/kg MTRN, p-value=0.0034 on day 14, Ptx + Vehicle vs. Ptx + 1.8 mg/kg MTRN, p-value=0.0008 on day 16, vehicle vs. Ptx + 0.5 mg/kg MTRN, p-value=0.0115, Ptx+ vehicle vs. Ptx + 1.8 mg/kg MTRN, p-value<0.0001 on day 18, vehicle vs. Ptx + 0.5 mg/kg MTRN, p-value=0.0179, Ptx + vehicle vs. Ptx + 1.8 mg/kg MTRN, p-value=0.0068 on day 20, vehicle vs. Ptx + 0.5 mg/kg MTRN, p-value= 0.0082, Ptx + vehicle vs. Ptx + 1.8 mg/kg MTRN, p-value=0.0010 on day 22, Ptx + vehicle vs. Ptx + 1.8 mg/kg MTRN, p-value=0.0435 on day 29, Ptx + vehicle vs. Ptx + 1.8 mg/kg MTRN, p-value=0.0225 on day 32, Ptx + vehicle vs. Ptx + 1.8 mg/kg MTRN, p-value=0.0244 on day 35). N=8/group until day 22 and N=4/group from day 26-57. C) Effect size calculated from the first dosage of MTRN injection (day1) till the end of testing shows a significant effect of rmMeteorin treatment compared to vehicle at both doses in paclitaxel treated mice (One-way ANOVA, F=5.361, p=0.0132, post-hoc Tukey, vehicle vs. Ptx + 0.5 mg/kg MTRN, p-value=0.0271, Ptx + vehicle vs. Ptx + 1.8 mg/kg MTRN, p-value=0.0245). Data represents mean ± SEM. Significance represented as Ptx + vehicle vs. 0.5 mg/kg MTRN* and Ptx + vehicle vs. 1.8 mg/kg MTRN#.

### 3.5 rmMeteorin reduced Satellite cell gliosis and gap junction expression in the prevention dosing paradigm in paclitaxel treated mice

We assessed whether administration of rmMeteorin attenuated satellite cell gliosis in the prevention dosing paradigm. We observed an increase in expression of GS in the paclitaxel treated cohort and this effect was significantly blocked in both groups treated with rmMeteorin (0.5 mg/kg and 1.8 mg/kg) (Fig 5A-C). Next, we examined the expression of connexin43 in the DRG in mice treated with the prevention dosing paradigm. We observed a significant reduction in connexin43 expression in animals treated with both doses of rmMeteorin (0.5 mg/kg or 1.8 mg/kg) compared to the paclitaxel treated cohort alone (Fig 5D, E).

**Figure 5:**
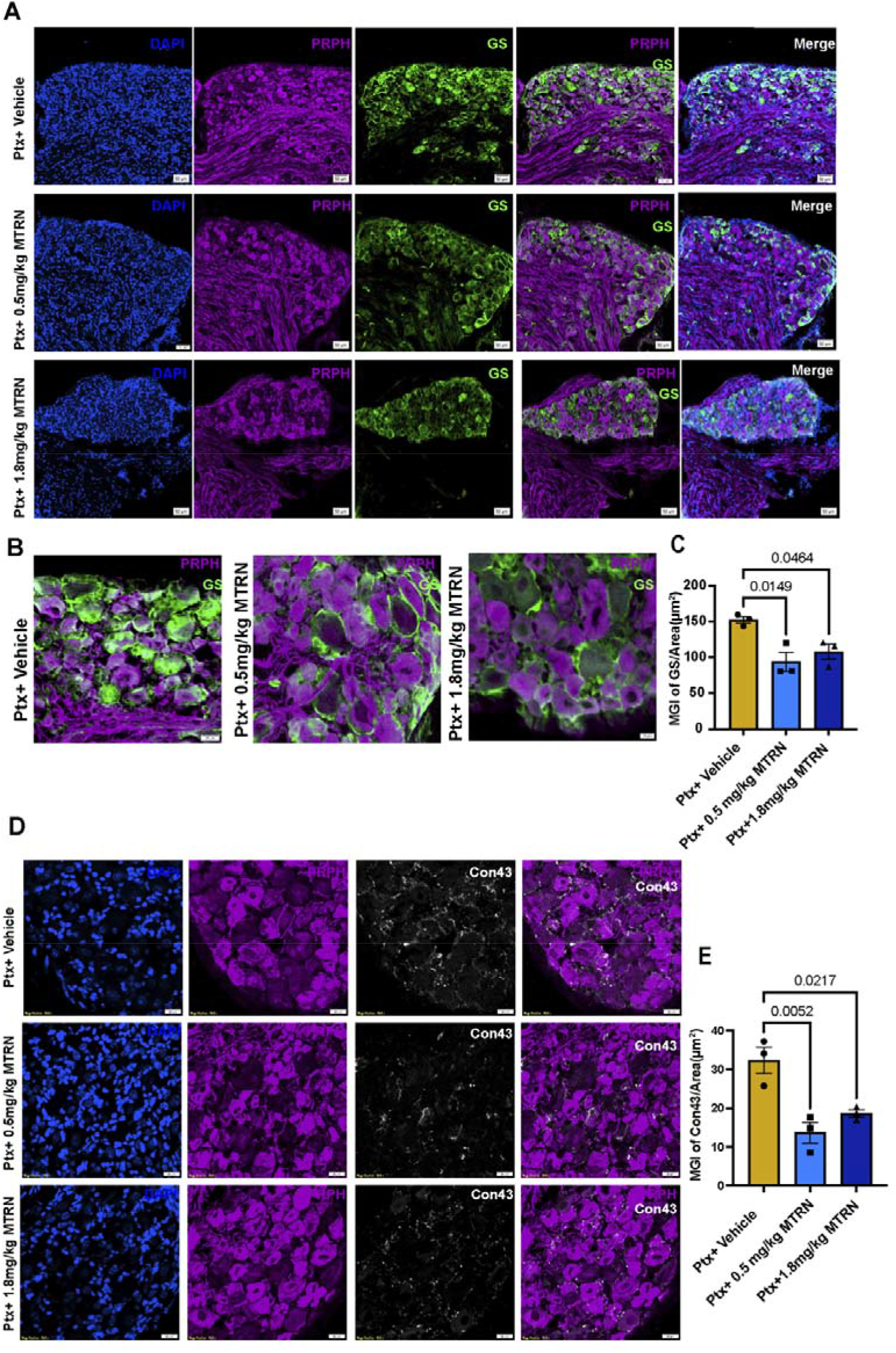
rmMeteorin reduces satellite cell gliosis and gap junction expression in the prevention dosing paradigm of paclitaxel treated mice. A-B) Representative image of satellite cell gliosis measured using GS immunoreactivity (Watkins et al.), peripherin (purple), and DAPI (blue). rmMeteorin reduced satellite cell gliosis in the DRG. C) Significant reduction in GS immunolabelling was observed in mice administered 0.5 mg/kg (royal blue) and 1.8 mg/kg (navy blue) MTRN in the prevention dosing paradigm compared to paclitaxel + vehicle (yellow) treated mice (One-way ANOVA, F=9.207, p-value=0.0148, post-hoc Tukey, Ptx + vehicle vs. Ptx + 0.5 mg/kg MTRN, p-value= 0.0149, Ptx + vehicle vs. Ptx + 1.8 mg/kg MTRN, p-value=0.0464). D) Representative images for the expression of gap junctions using Connexin 43 immunolabeling (white), peripherin (purple), DAPI (blue). E) Decreased expression of connexin 43 immunoreactivity was observed in mice treated with rmMeteorin (One-way ANOVA, F=14.12, p-value=0.0054, post-hoc Tukey, Ptx + vehicle vs. Ptx + 0.5 mg/kg MTRN, p-value=0.0052, Ptx + vehicle vs. Ptx + 1.8 mg/kg MTRN, p-value=0.0217) N = 3/group. Data are represented as mean ± SEMs. Scale bar = 50μm (A), 20μm (B and D) and 10μm (B, far right panel).

### 3.6: rmMeteorin partially reverses IENF density loss caused by paclitaxel

Loss of IENFs is a characteristic sign of chemotherapy-induced neuropathy. Paclitaxel treatment is known to cause the retraction of nerve fibers from the epidermis in animals and in patients (Boyette-Davis et al., 2011). We assessed the density of IENFs using PGP9.5 immunostaining. Paclitaxel treatment induced significant loss of IENF in the skin compared to naïve mice. Administration of both doses of rmMeteorin (0.5mg/kg or 1.8mg/kg) in the prevention dosing paradigm partially reversed the IENF loss caused by paclitaxel (Fig 6 A, B).

**Figure 6:**
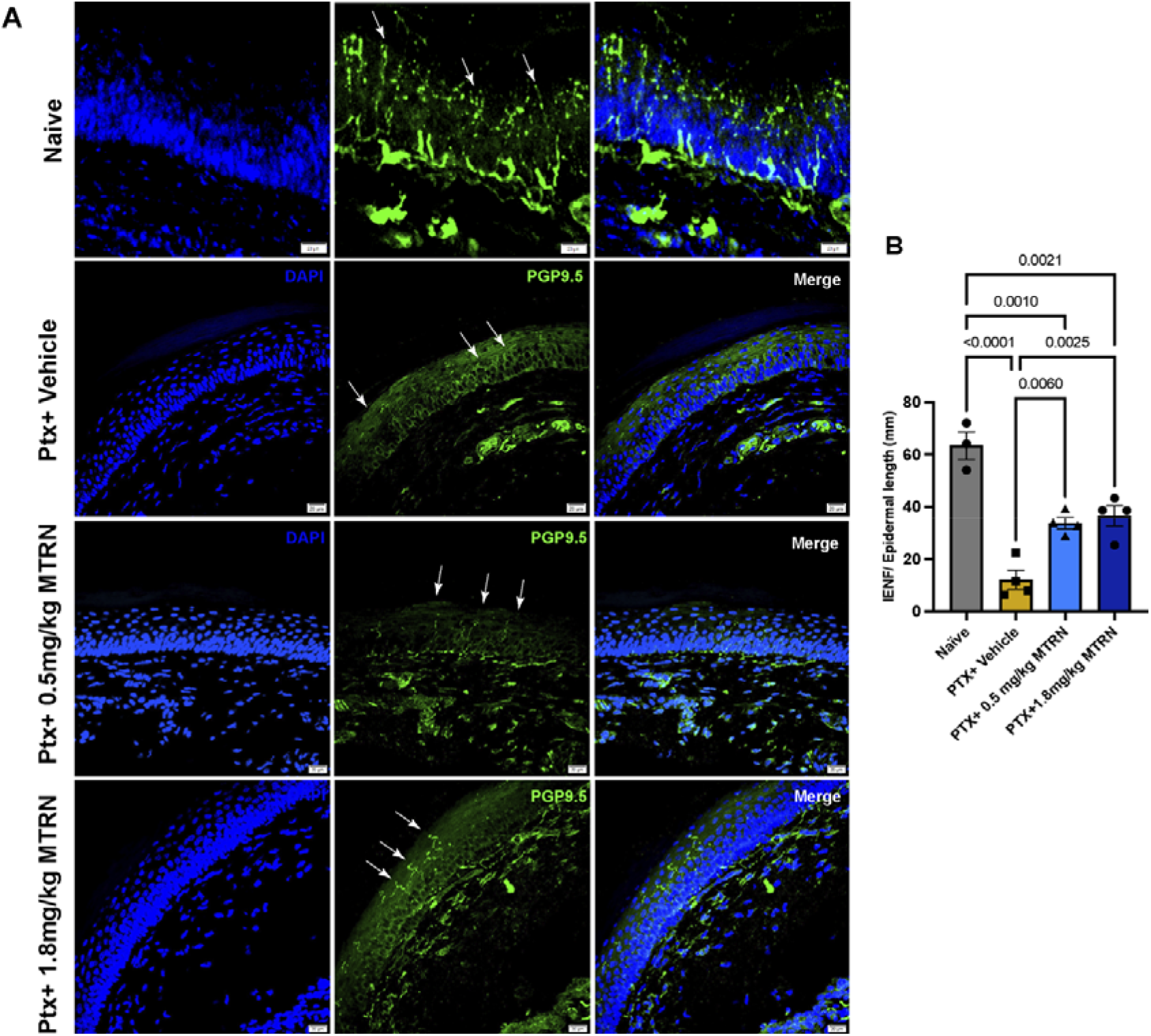
Systemic administration of rmMeteorin partially reverses IENF density loss caused by paclitaxel in the prevention dosing paradigm. A) Representative image of skin from hind paw immunolabeled for fibers with PGP9.5 (Watkins et al.) and nuclei with DAPI (blue). B) rmMeteorin administration (0.5 mg/kg and 1.8 mg/kg) partially reversed the IENF density loss caused by paclitaxel. Ptx + 0.5 mg/kg MTRN (royal blue) and 1.8 mg/kg MTRN (navy blue) showed significant increased nerve fiber density compared to the Ptx + vehicle group (One-way ANOVA, F=29.68, p-value<0.0001, post-hoc Tukey, Naïve vs. Ptx + vehicle, p-value<0.0001, naïve vs. Ptx + 0.5mg/kg MTRN, p-value=0.0010, naïve vs. Ptx + 1.8 mg/kg MTRN, p-value= 0.0021, Ptx + vehicle vs. Ptx + 0.5 mg/kg MTRN, p-value=0.006, Ptx + vehicle vs. Ptx + 1.8 mg/kg, p-value=0.0025) N = 3/group. Data are represented as mean ± SEMs. Scale bar = 20μm.

## Discussion

In this study, we demonstrated that systemic administration of rmMeteorin results in the resolution of mechanical hypersensitivity in paclitaxel-induced peripheral neuropathy in male and female mice. Our findings align with the previous demonstration that Meteorin promotes an antinociceptive effect in peripheral neuropathy models where the neuropathic pain is caused by traumatic injury to peripheral nerves (Jorgensen et al., 2012; Xie et al., 2019b). Our findings also demonstrate that rmMeteorin treatment reverses two signs of satellite cell gliosis, consisting of increased GS expression and enhanced expression of the gap junction protein connexin 43. Finally, rmMeteorin partially prevented the retraction of IENFs from the skin of mice treated with paclitaxel. Collectively, these experiments demonstrate that rmMeteorin has a beneficial effect on multiple aspects of paclitaxel-induced neuropathy including improved behavioral and molecular and cellular outcomes.

Recently, KIT receptor tyrosine kinase has been identified in cardiac tissue as a putative receptor for the sibling protein Meterorin-like (Reboll et al., 2022). Although, the receptor-mediated mechanism of action of Meteorin remains unknown, during development meteorin is known to play an important role in differentiation of glial cells, including satellite glial cells in the DRG (Nishino et al., 2004). Reactive gliosis is characterized by morphological and molecular changes in peripheral and/or central glial cells and is caused by injury to peripheral nerves by trauma or by chemotherapeutic treatment (Nishino et al., 2004; Lee et al., 2015; Ji et al., 2016) Satellite cell and astrocytic gliosis results in upregulation of glial fibrillary acidic protein and GS, cellular hypertrophy, and proliferation. In the adult DRG, reactive gliosis caused by Transforming Growth Factor-ß1 (TGF-ß1) triggers the production of endogenous Meteorin, resulting in negative feedback loop promoting resolution of gliosis (Lee et al., 2015). The existing literature suggests a clear role for Meteorin-mediated effects on satellite glial cells but developmental signaling effects may be quite different than those observed on adult cells, which is common for growth factors. Our results show that administration of rmMeteorin reversed satellite cell gliosis after paclitaxel treatment in both reversal and prevention dosing paradigms. While more work is needed to understand the underlying mechanism, our findings support a potential role for Meteorin in disease-modifying effects with respect to satellite glial cells in paclitaxel-induced neuropathy.

Satellite glial cells tightly envelop neurons in the DRG (Hanani et al., 2002; Hanani, 2005; Huang et al., 2010; Warwick and Hanani, 2013). Gap junction-mediated coupling between satellite glial cells is increased after administration of chemotherapy drugs such as paclitaxel and oxaliplatin and this coupling potentially contributes to pain in CIPN (Warwick and Hanani, 2013). Our results are consistent with the existing literature where paclitaxel induces an increase in the expression of connexin 43 in rodent DRG in satellite glial cells (Warwick and Hanani, 2013). In our experiments, administration of rmMeteorin reduced the expression of connexin 43 in mouse DRG. Previous studies demonstrate that reducing connexin 43 expression in neuropathic pain models in rodents results in diminished pain behaviors, consistent with the notion that reducing satellite glial cell coupling is beneficial for neuropathic pain (Hanani, 2005; Ohara et al., 2009; Warwick and Hanani, 2013; Amatore et al., 2020). While we observed clear effects of rmMeteorin in the reversal dosing paradigm at the lower dose in male and female mice, effects at the higher dose were not significant in female mice. In the prevention dosing paradigm, significant effects were seen in females at both doses on satellite glial cell markers. The reason for this small sex difference at the higher dose of rmMeteorin in the reversal dosing paradigm is not clear at this time.

Previous studies have shown that Meteorin causes extensive neurite growth in small-and intermediate sized neurons of the developing sensory ganglion, including nociceptors (Nishino et al., 2004). Meteorin treatment also promotes neurite outgrowth from DRG explants of neonatal animals (Nishino et al., 2004). This could suggest that in addition to an effect on satellite glial cells, Meteorin may also have a direct action on DRG neurons. Consistent with preclinical and clinical observations with paclitaxel, we observed a loss in IENF density in the skin following chemotherapy treatment. this IENF loss was partially reversed in the prevention dosing paradigm by rmMeteorin at both doses. A shortcoming of this experiment is that it was done only in females, but the greater impact of paclitaxel neurotoxicity on females justifies this choice (Ozdemir et al., 2018). While we favor the hypothesis that rmMeteorin acts directly on nerve endings to protect them from paclitaxel toxicity, it is also possible that Meteorin acts on glial cells surrounding the neuron, to promote breakdown of neurotoxic metabolites, and releases neuroprotective factors that promote the survival of neuronal endings in the epidermis (Lee et al., 2010; Lee et al., 2015).

There are some limitations to our study. Notably, the complete mechanism of action through which rmMeteorin reduces paclitaxel-induced peripheral neuropathy is not resolved by our work. This will be a primary goal of future studies. Moreover, this study and others in the literature (Jorgensen et al., 2012; Xie et al., 2019a) have only assessed the effect of Meteorin in rodent models and on rodent cells. Bulk, spatial and single-cell sequencing experiments suggest that the *METRN* gene is highly expressed by neurons and likely glial cells in human DRG, consistent with similar experiments in mice (Ray et al., 2018; Zeisel et al., 2018; Tavares-Ferreira et al., 2022). A key future direction will be to conduct validation studies in human DRG using recombinant human Meteorin to create a compelling rationale for further clinical development. Finally, as noted above, we did not conduct all studies in both sexes, but we believe that the findings from the reversal dosing paradigm strongly suggest that consistent effects should be observed. Examining potential differences in Meteorin signaling on male and female human DRG cells will be a critical step to understanding the future potential of this therapeutic approach in clinic.

## Notes

### Competing Interest Statement

Torsten M. Madsen, Kenneth A. Petersen, and Gordon Munro are employees of Hoba Therapeutics.

